# Metabolism Control in 3D Printed Living Materials

**DOI:** 10.1101/2021.01.15.426505

**Authors:** Tobias Butelmann, Hans Priks, Zoel Parent, Trevor G. Johnston, Tarmo Tamm, Alshakim Nelson, Petri-Jaan Lahtvee, Rahul Kumar

**Affiliations:** Institute of Technology, University of Tartu, Tartu 50411, Estonia; Institute for Macromolecular Chemistry, University of Freiburg, Freiburg 79104, Germany; Department of Chemistry, University of Washington, Seattle, WA 98195, USA

**Keywords:** living materials, hydrogel, 3D printing, yeast, cell isolation, cell size, flow cytometry, oxygen, macromolecules, metabolism, beer fermentation, biotechnology

## Abstract

The three-dimensional printing of cells offers an attractive opportunity to design and develop innovative biotechnological applications, such as the fabrication of biosensors or modular bioreactors. Living materials (LMs) are cross-linked polymeric hydrogel matrices containing cells, and recently, one of the most deployed LMs consists of F127-bis-urethane methacrylate (F127-BUM). The material properties of F127-BUM allow reproducible 3D printing and stability of LMs in physiological environments. These materials are permissible for small molecules like glucose and ethanol. However, no information is available for oxygen, which is essential— for example, towards the development of aerobic bioprocesses using microbial cell factories. To address this challenge, we investigated the role of oxygen as a terminal electron acceptor in the budding yeast’s respiratory chain and determined its permissibility in LMs. We quantified the ability of cell-retaining LMs to utilize oxygen and compared it with cells in suspension culture. We found that the cells’ ability to consume oxygen was heavily impaired inside LMs, indicating that the metabolism mostly relied on fermentation instead of respiration. To demonstrate an application of these 3D printed LMs, we evaluated a comparative brewing process. The analysis showed a significantly higher (3.7%) ethanol production using 3D printed LMs than traditional brewing, indicating an efficient control of the metabolism. Towards molecular and systems biology studies using LMs, we developed a highly reliable method to isolate cells from LMs for flow cytometry and further purified macromolecules (proteins, RNA, and DNA). Our results show the application of F127-BUM-based LMs for microaerobic processes and envision the development of diverse bioprocesses using versatile LMs in the future.

## Introduction

The application of three-dimensional (3D) printing or additive manufacturing is a rapidly advancing field in biology after it has first been innovated and demonstrated in applied materials science.^1,2^ Initially, most of the bioprinting research has been carried out in the domain of tissue engineering, from where it has advanced to the development of modular bioreactors.^3–8^ However, more recently, applications of this technology are demonstrated in microbial 3D printing for investigating infections using biofilms, bioremediation, on-demand biomanufacturing, and biosensors using so-called living materials (LMs).^6,9–13^ The triblock copolymer Pluronic F127, is extensively applied in direct-write or extrusion 3D printing, constituting a common bioink in both tissue engineering and biotechnology.^14–20^ This versatile polymer shows stimuli-responsive properties, which among others are triggered in response to temperature and pressure changes.^21^ Such properties allow a homogenous mixing of cells and other materials and enable 3D printing. Additionally, the modification of the polymer chain ends facilitates their crosslinking in a photoinitiated reaction leading to tough and resilient materials.^22^ F127-based LMs are already applied in biotechnological processes using yeast, modular biomanufacturing using co-culture systems, and the design of multi-kingdom microorganismal networks for developing modular bioreactors for chemicals production.^17,19,23^ Such LMs have immense potential for emergent biotechnological and medical applications, among which biosensors and efficient biomanufacturing platforms can lead to transformative technologies and products.

Mammalian cells generally require oxygen to thrive, but their encapsulation inside of a 3D tissue culture matrix impairs the sufficient oxygen supply.^24,25^ Not only for mammalian cells, but also for microbes— oxygen plays a pivotal role. Some microorganisms are strictly anaerobic, whereas others are facultative aerobes.^26,27^ The latter’s metabolic adaptability to the presence/absence of oxygen allows their use in biotechnology to produce e.g., ethanol or other biomolecules.^28^ In aerobic organisms, oxygen functions as a terminal electron acceptor and allows a highly efficient cellular energy generation.^29^ The compartmentalization of some metabolic pathways in mitochondria, the site of oxygen consumption, allows a productivity increase by 260% for commercially important branched-chain alcohols (isobutanol) and highlights the need for designing efficient oxygen diffusion in LMs.^30^ However, anaerobic organisms such as acetogens or anammox bacteria do not require oxygen, but instead have evolved to use alternative substrates assimilation and cofactors regeneration strategies to produce energy and biomass, and oxygen impermissible LMs could be applicable for cultivating such microbes.^31–33^

Despite the advances demonstrating possible processes and products, a better understanding of LMs is yet necessary to utilize the cellular capabilities attained upon encapsulation. Further, comprehending cellular changes inside LMs can enable the design of tailored bioprocesses and applications. Additionally, in most of the studies so far, cells are often not fully retained in LMs during the cultivation, making it challenging to compare the results between suspension and immobilized cells. This highlights the importance of investigating the encapsulated cells in cell-retaining LMs, allowing the performance comparison and process optimization relative to suspension cultures. Nagarajan and colleagues showed the decoupling of the cellular metabolism from growth in calcium-alginate immobilized cells, despite only partial cell-retention.^34^ That study highlights the importance of creating and understanding stable cell-retaining LMs since calcium-alginate systems tend to be unstable. Cell-retaining LMs could significantly enhance process productivity by directing carbon towards products instead of the formation of biomass. Previously, we showed that the encapsulated cells exhibit an altered cellular size, but information on their metabolism is still lacking.^20^

In this study, we investigated 3D printed living materials composed of the budding yeast *Saccharomyces cerevisiae* and the polymer F127-bis-urethane methacrylate, previously deployed for the encapsulation of microorganisms.^19,20,23^ Figure 1 presents a detailed overview of our study design. For the first time, we report cellular oxygen utilization in F127-based yeast-laden cell-retaining hydrogels and the metabolic mode of the encapsulated yeast. Further, we developed a reliable method for isolating cells from LMs after cultivation, applied flow cytometry, and purified macromolecules, providing a valuable resource for the characterization of these LMs in the future. We used the metabolic mode information for the demonstration of a more efficient beer fermentation.

**Figure 1:**
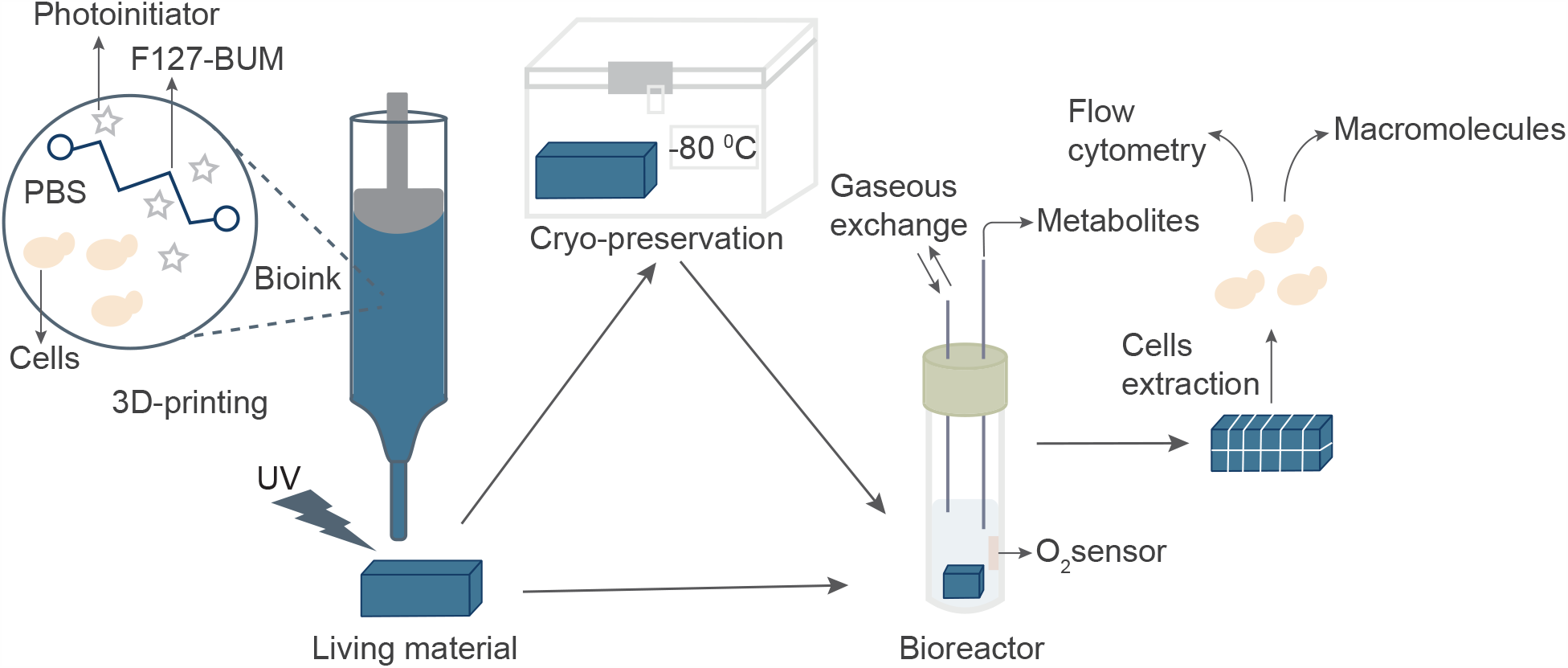
Workflow of the experiments. 3D printing of a bioink consisting of PBS, F127-BUM, yeast cells and a photoinitiator for UV curation of LMs; direct cultivation or freeze-thaw cultivation (after storage in −80°C) with oxygen and HPLC measurements; post-cultivation isolation of cells with flow cytometry analysis and macromolecule purification.

## Materials and Methods

### Chemical Synthesis of Pluronic F127-BUM

An F127-derived polymer, namely F127-bis-urethane methacrylate, was provided by the Nelson laboratory at the University of Washington. Millik and colleagues reported the details of the polymer synthesis earlier.^18^ Briefly, pluronic F127 was dried and subsequently dissolved in anhydrous dichloromethane (DCM). This was followed by the addition of dibutyltin dilaurate to the mixture and followed by the dropwise addition of a 2-isocyanatoethyl methacrylate/DCM solution. The reaction was stirred for 2 d, quenched with methanol, and precipitated in diethyl ether. The polymer was collected via centrifuge and washed twice with fresh ether. A fluffy white powder texture of the polymer was achieved by drying it under vacuum.

### Hydrogel Preparation for 3D Printing

Sterile phosphate-buffered saline (PBS) was mixed with F127-BUM and cooled at 4 °C overnight to prepare a 30 wt % solution. One liter of PBS (pH = 7.2) contained 8 g of NaCl (Sigma-Aldrich),1.44 g of Na_2_HPO_4_ (Fisher Scientific), 0.24 g of KH_2_PO_4_ and 0.2 g of KCl in milli-Q water. To make a hydrogel ready for printing, 1.5 µL g^-1^ solution of the photoinitiator 2-hydroxy-2-methyl propiophenone (Irgacure 1173; >97 %, Sigma-Aldrich) was added at 4°C temperature. Spun-down cells (10^6^ cells g^-1^ solution) were added to the solution. A short stirring of both additives ensured uniform distribution. After incubating 30 min on ice, to make the mixture bubble-free, it was poured into a 10 mL dispensing barrel equipped with a 0.41 mm dispensing tip (both manufactured by Adhesive Dispensing, United Kingdom) and warmed to room temperature to transform to a shear-responsive state for printing. We used 10^7^ cells g^-1^ solution for the scaled-up demonstration of beer fermentation.

### 3D Printing

3D printing was performed on a K8200 printer (Velleman, Belgium) modified to be applicable for direct pressure dispensation. The computer-aided design models were fabricated with SketchUp Make 2017 (Trimble, USA), the printer was controlled with Repetier-Host (Hot-World, Germany) and the G-code was generated using the 3D printing toolbox Slic3r. The model’s dimensions were 10 × 3 × 3.5 mm (X, Y, Z), sliced with one outer perimeter and printing was performed in the vase mode with a print speed of 10 mm s^-1^. Directly after the print, the hydrogel was cross-linked for 60 s with four light-emitting diodes (CUN66A1B, Seoul Viosys, Republic of Korea) emitting at a wavelength of 365-367 nm. Afterwards, the living materials were washed in 70 % ethanol for 60 s to avoid viable yeast or any potential microbial contaminants on the surface of living materials, allowing a cultivation solely based on the cells immobilized inside of the hydrogel.

For the scaled-up pilot demonstration of beer fermentation, a model with 50 × 7 × 7 mm (X, Y, Z) was used, sliced with one outer perimeter, 45 % grid infill, varying print speed for perimeters and infill to fabricate the structure. UV curation was extended to 120 s while other steps were followed as described above before initiating the brewing process.

### Yeast Strain, Media and Cultivation Conditions

The yeast strain *Saccharomyces cerevisiae* CEN.PK113-7D (*MATa, MAL2-8*^*c*^, *SUC2*) was used in the study and cultivated in chemically defined minimal medium (MM). The beer yeast Bry-97 (Lallemand, Canada) was used for the alcohol fermentation of wort. The composition of 1 L MM1 (pH = 6) was 10 g glucose (Acros Organics), 5 g of (NH_4_)_2_SO_4_ (Lach-Ner), 3 g of KH_2_PO_4_ (Sigma-Aldrich) and 0.5 g of MgSO_4_ ·7 H_2_O (Sigma-Aldrich) in milli-Q water. The composition of 1 L MM2 (pH = 6.9) was 5 g glucose (Acros Organics), 2.5 g of (NH_4_)_2_SO_4_ (Lach-Ner), 3 g of KH_2_PO_4_ (Sigma-Aldrich), 5.25 g of K_2_HPO_4_ (Merck) and 0.25 g of MgSO_4_ (Sigma-Aldrich) in milli-Q water. One milliliter trace elements (all Sigma-Aldrich, unless marked differently) and 1 mL vitamin solution (all Sigma-Aldrich, unless marked differently) were added after sterilization of both MM. One liter trace element solution (pH = 4) contained EDTA sodium salt (Lach-Ner), 15.0 g; ZnSO_4_·7H_2_O, 4.5 g; MnCl_2_·2 H_2_O, 0.84 g; CoCl_2_·6 H_2_O, 0.3 g; CuSO_4_·5 H_2_O, 0.3 g; Na_2_MoO_4_·2 H_2_O, 0.4 g; CaCl_2_·2 H_2_O (Carl Roth), 4.5 g; FeSO_4_·7 H_2_O, 3.0 g; H_3_BO_3_, 1.0 g; and KI, 0.10 g. One liter vitamin solution (pH = 6.5) contained biotin, 0.05 g; p-amino benzoic acid, 0.2 g; nicotinic acid, 1 g; Ca-pantothenate, 1 g; pyridoxine-HCl, 1 g; thiamine-HCl, 1 g; and myoinositol (AppliChem), 25 g; in milli-Q water. The mash contained 200 g/L Best Pale Ale malt (Simpsons, United Kingdom), and the wort was supplemented with 3.6 g Columbus hops (BarthHaas, Germany), resulting in 1 L of wort.

After 3D printing and washing in ethanol, the LMs were cultivated for 16 h in 5 mL MM1. Subsequently, they were cryopreserved in 20 % glycerol 1:2 MM1 at −80 °C to use LMs fabricated from one batch and printing session.

After mildly thawing (−20 °C, on ice, 5 °C and room temperature, each 1 h) and washing LMs or suspension cells with PBS, the second or direct cultivation was carried out at 30 °C for the given times at 150 rpm in 10 mL culture medium with either glucose (MM2) or ethanol (MM3) as a carbon source in 50 mL sterilized glass tubes (D2 Biotech, Sweden) equipped with filters.

LMs with Bry-97 were ethanol-treated, washed in milli-Q water, and directly used in the experiment for 14 d at 20 °C in 180 mL glass bottles, equipped with airlocks.

### Dissolved Oxygen

The oxygen data were acquired with a PICFIB2 fiber and OXSP5 sensor spots (both PyroScience, Germany), which were attached to the cultivation tubes. Since *S. cerevisiae* can consume ethanol in the presence of oxygen, it was used in the culture medium (MM3) to cross-validate dissolved oxygen data obtained in the presence of glucose as a carbon source (MM2).

### High-Performance Liquid Chromatography

High-performance liquid chromatography (HPLC) was performed using a Rezex ROA-Organic Acid column (Phenomenex, USA) with 5 mM sulfuric acid (>99.5 %, Merck) as a mobile phase at 45 °C for ethanol, glycerol and acetic acid measurements and a Rezex RPM-Monosaccharide column (Phenomenex, USA) at 80 °C with milli-Q water as mobile phase for sugars. A Shimadzu Prominence-i LC-2030C Plus (Japan) equipped with a Refractive Index Detector RID-20A (Shimadzu, Japan) was used for analysis from filtered culture samples.

### LM and Cell Mass, Determination of Cell Numbers

At the given points in time, an LM was taken out from the medium (30 °C) and weighed on a precision scale (Adam Equipment, UK). To determine the dry weight of the cells, the suspension was filtered over a 0.45 μm membrane and washed with distilled water, dried at 65 °C overnight and weighed on the precision scale.

For cell counting, 10 µL of a cell suspension was pipetted into a Neubauer cell counting chamber. Cells were counted and the total amount of cells calculated. If necessary, a cell suspension sample was mixed 1:1 with 0.4% trypan blue and the viability was assessed in the counting chamber.

To estimate the total number of cells per LM structure, the following assumptions and calculations were considered. The average print weight was taken to calculate the cell inoculum per structure. Due to the ethanol treatment and based on the published data, it was estimated that approximately 10 % of the cells were killed.^20^ The structures were incubated for 24 h, weighed afterwards and cut into thin slices with a scalpel to measure cell colony diameters under the microscope. The average cell volume, as well as the colony volume and weight increase over time, were taken into consideration for the calculation of the total number of cells in a living material after the cultivation.^20^ ImageJ 1.52i was used to measure the diameter of 40 cell colonies in a sliced sample after the mentioned incubation.

An Eclipse Ci-L (Nikon, Japan) microscope was used for imaging with common microscope glass slides and cover glasses.

### Cell Isolation from LMs

For capturing a snapshot of the cellular and macromolecular landscape during the cultivation— freezing of cultivated cells in liquid nitrogen is a commonly used method, and here, it was applied for LMs. After cultivation, the LMs were quenched in liquid nitrogen. The samples were cut in small pieces with a scalpel, washed with PBS, vortexed, centrifuged and resuspended, and finally filtered (pore size 20 µm) to recover the cells in the filtrate. At least 100,000 cells were further collected for flow cytometry. The remaining LM debris was treated in a FastPrep-24 homogenizer (MP Biomedicals), 4 m s^-1^ and 20 s cycle to isolate more cells.

Suspension cells were pelleted at 3200 rpm for 3 min, supernatant was decanted, and the pellet-containing tube was frozen in liquid nitrogen and stored at −80 °C until taken for further analysis or purification.

### Flow Cytometry

Isolated cells recovered after cutting, or processed suspension cells were washed with PBS and fixed with ice-cold ethanol (final concentration 70%) to store the cells at 4 °C for a maximum of one week. Fixed cells were centrifuged and carefully resuspended in PBS for flow cytometry analysis. Flow cytometry analysis was carried out using an Attune NxT Acoustic Focusing Cytometer (Invitrogen, USA). The flow cytometry data was analyzed using the FlowJo software (BD Biosciences, USA). At least 100,000 events were collected for every sample. Standard beads with a 6 μm diameter were used as a reference along with every sample (10,000 events).

### Macromolecule Purification and PCR

Genomic DNA was purified according to the protocol by Lõoke and colleagues.^35^ For RNA purification, the cells were processed using the YeaStar RNA Kit (Zymo Research, USA) according to the manufacturer’s protocol. Total proteome was obtained using the Y-PER reagent with yeast cells in a glass beads-containing tube and by disruption of the cells in a FastPrep-24 homogenizer (MP Biomedicals, USA). For cell disruption, 4 m s^-1^ and 20 s cycles were applied for 10 times with 5 min intervals between each cycle.

The purified molecules were quantified using a NanoDrop ND-1000 spectrophotometer (Thermo Fisher Scientific, USA).

A volume of 1 µL of the purified DNA was taken as a template for amplification of the *URA3* gene with primers 1 and 2. The reverse transcription and following polymerase chain reaction (PCR) was carried out using the RevertAid First Strand cDNA Synthesis kit (Thermo Scientific) and primers NL-1 and NL-4 (Table 1).

**Table 1:**
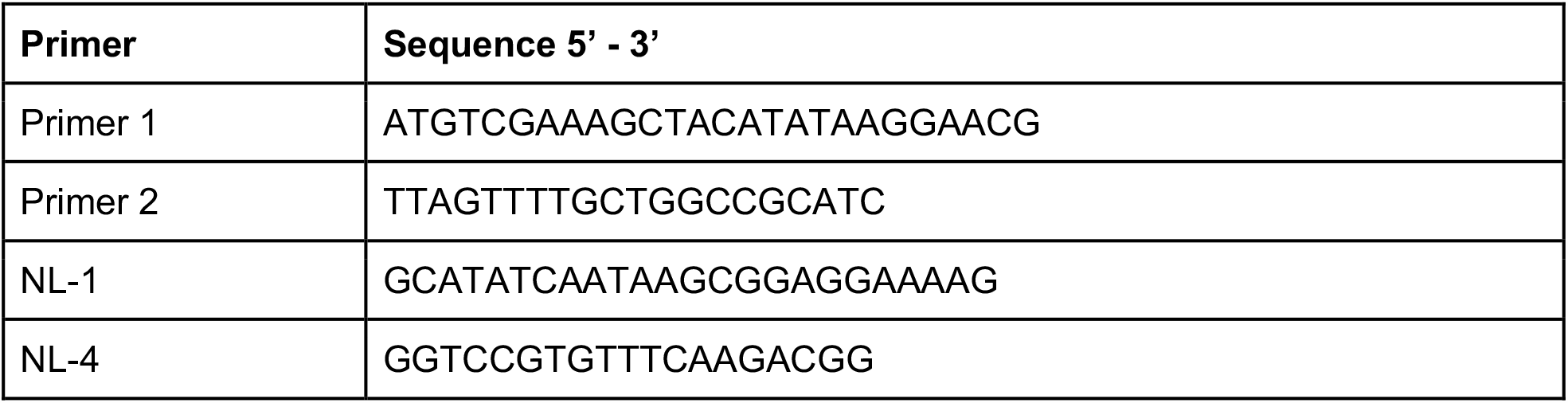
Primer sequences used for the *URA3* gene (Primer 1, 2) and D1/D2 rRNA (NL-1,4).

Polymerase chain reactions were performed with 35 cycles of 30 s denaturation, 30 s annealing and elongation depending on the fragment length using the DreamTaq Green PCR Master Mix (Thermo Scientific, Lithuania). The GeneRuler 1 kb Plus ladder (Thermo Scientific, Lithuania) was used as a marker.

## Results and Discussion

In this study, we compared LMs that were freshly 3D printed (FP) or 3D printed and cryopreserved (CP) and benchmarked their performance against that of suspension cells (SCs). We printed and cultivated the budding yeast *S. cerevisiae* CEN.PK113-7D as LMs for 16 h in MM1 as pre-culture. The experiments followed two different cultivation strategies: (i) print-perform (PP) using FP–LMs, and (ii) print-preserve-perform (PPP) using CP–LMs. Both were cultivated in MM2 for 40 h in the perform stage. The latter strategy was tested to investigate the storage and re-usage potential of LMs that could make the cultivation process more efficient and scalable by avoiding a fresh batch printing per cultivation. For this, we cryopreserved the fabricated LMs in 20 % glycerol containing MM1 at −80 °C. To ensure a direct comparison of both FP and CP–LMs with SCs, the latter were either directly cultivated from pre-cultures (SCs) or taken from similarly cryopreserved cultures (CP–SCs). This way, we ensured experimentation using CP–SCs and CP-LMs originated from a single cryopreserved batch and demonstrated the usability of cryopreserved LMs. If not indicated otherwise, those samples from the cryopreserved batch were used throughout this study. The inoculum of SCs and CP–SCs was equal to the cells per LMs and contained 1.5 − 2.0 × 10^7^ cells. An assessment of cellular viability of thawed CP samples allowed us to show that the cells in CP-LMs were equally viable (48.6 ±2.9 %) as in CP-SCs (45.4 ±0.6 %) (Figure S1), indicating the suitability of CP-LMs for functioning as inoculum for initiating experiments. We collected the present data using fully cell-retaining LMs, allowing us to contrast SC and LM cultures.

### omparative Physiology of CP–SCs and CP–LMs

We compared the yeast physiology as exhibited by CP–SCs and CP–LMs in the presence of glucose (in MM2) by measuring extracellular metabolites and dissolved oxygen (Figure 2, Table 2). Budding yeast produces ethanol, glycerol, acetate and CO_2_ as metabolic byproducts when using glucose as a carbon source in batch cultures. To obtain physiologically comparable data, we cultivated both CP–SCs and CP–LMs during their glucose uptake in independent triplicate experiments to assess their metabolic performances. We found that glucose was consumed faster by CP–SCs than CP–LMs, where the glucose-phase almost ended in 16 h for CP–SCs (Figure 2A), but it took longer than 40 h for completion of the glucose phase for CP–LMs (Figure 2B). Additionally, both glucose and oxygen consumption patterns differed between CP–SCs and CP– LMs. The glucose consumption pattern in CP–LMs appeared to be linear rather than exponential (Figure 2), highlighting the impact of physical confinement in a hydrogel matrix and a potential diffusion limitation for substrates or metabolites. Moreover, compared with glucose, the oxygen consumption, estimated based on dissolved oxygen profile, was almost negligible in CP–LMs. It appeared that yeast primarily consumed glucose by the fermentative metabolism in CP–LMs and the respiratory chain in mitochondria, which uses oxygen as a terminal electron acceptor in aerobic cultivation, did not have access to oxygen (Figure 2B, Table 2). To rule out the possibility that this negative impact on oxygen consumption was not due to the cryopreservation of LMs, we cultivated FP-LMs under the same conditions. We found a similar physiology pattern as CP-LMs (Figure S2, Table ST1). These observations imply that the budding yeast’s physiology observed in CP-LMs was not due to cryopreservation but due to the physical confinement in the F127-BUM hydrogel. Our results therefore suggest that the PPP strategy is deployable for cultivating LMs.

**Table 2.** Yields on glucose (G). Calculation done over the complete time span. ^1*^ *p* < 0.05 (one tail), ^**^ *p* < 0.01 significant difference from suspension cells, n = 3.

| | $Y_{G, \text{glycerol}} \text{ (g/g)}$ | $Y_{G, \text{acetate}} \text{ (g/g)}$ | $Y_{G, \text{ethanol}} \text{ (g/g)}$ |
| --- | --- | --- | --- |
| CP-SCs (Mean ± Std) | 0.1013 ± 0.05 | 0.0204 ± 0.09 | 0.3927 ± 0.22 |
| CP-LMs (Mean ± Std) | 0.0932 ± 0.00 *<sup>1</sup> | 0.0180 ± 0.00 ** | 0.4459 ± 0.00 ** |

**Figure 2:**
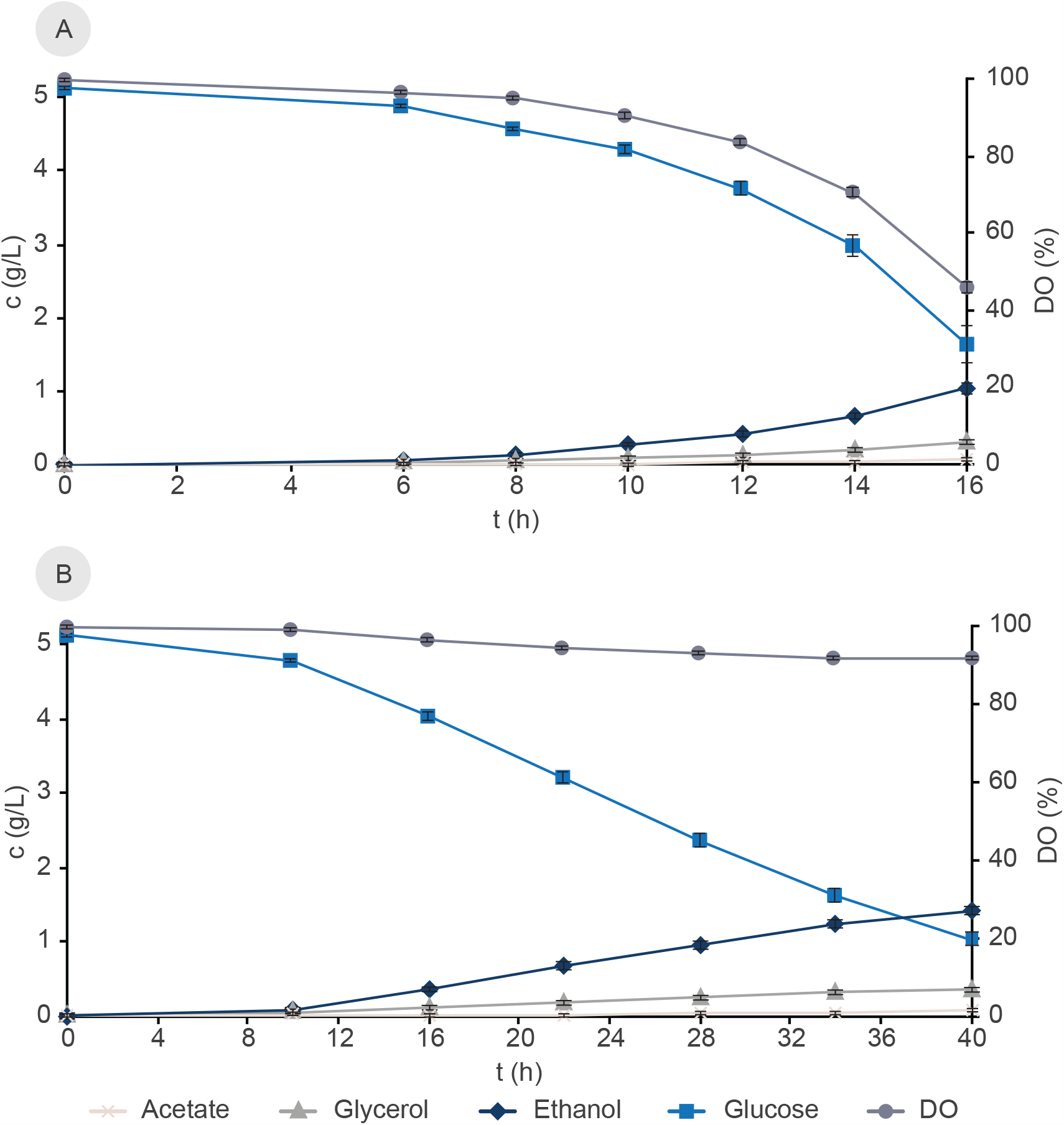
Yeast physiology in glucose phases of batch cultivations. (A) CP-SCs (B) CP-LMs. DO: dissolved oxygen. Error bars represent standard deviations, n = 3.

The byproduct yields during the batch cultivation for CP–LMs were significantly different from CP-SCs (Table 2). Notably, the ethanol yield on glucose was 13.6 % higher in CP–LMs than CP-SCs (Table 2). The dry weight of a cell from suspension culture (CP-SC: 24.5 pg) was slightly higher than previously reported numbers, while cells extracted from LMs appeared to be lighter compared to SCs (CP-LM: 12.7 pg) likely due to changes in the cellular phenotype.^20,36,37^

### Oxygen limitation in LMs

Since glucose-grown LMs did not show a typical oxygen consumption pattern based on the dissolved oxygen measurements, we were interested in validating whether the immobilized cells could access oxygen properly. We reasoned that cultivating LMs in ethanol containing culture medium (MM3), which requires oxygen for complete consumption by yeast, could validate our findings on atypical oxygen consumption by LMs in glucose-grown cultures. For this, we initially pre-cultured both CP–SCs and CP–LMs in glucose-containing medium (MM2) for 16 h, providing enough cells for oxygen consumption and ruling out the possibility of the absence of oxygen consumption due to lack of a sufficient number of cells in LMs. After 16 h, we transferred both CP–SCs and CP–LMs to fresh culture medium (MM3), where glucose was replaced by ethanol as the primary carbon source (Figure 3). We found that CP–SCs expectedly consumed ethanol and showed biomass formation (Figure 3A). In contrast, CP–LMs failed to utilize ethanol to the same extent; the apparent decrease in ethanol level was likely due to its conversion to acetate (Figure 3B). Ethanol to acetate conversion does not require oxygen. We also considered the ethanol evaporation by subtracting the final evaporated ethanol from a control MM3 without cells and calculated that only about 5.7 % of ethanol converted to acetate by CP–LMs after 180 h of cultivation (Figure 3B). After finishing the ethanol phase experiments, we surprisingly noticed a disproportionate increase in CP–LMs mass and cell size therein as measured by flow cytometry (Figure S3). To investigate the reason for this observation, we microscopically examined cells isolated from CP–LMs and noticed an increased vacuole size (Figure S4). This is a typical response to environmental stress in cell cultures.^38,39^ Usually, one would expect a decreased cell size during ethanol-phase growth in suspension cultures.^40^ We also observed a decrease in cell viability (93.54 %) after cultivation on ethanol as a carbon source, indicating cellular stress (Figure S4B).

**Figure 3:**
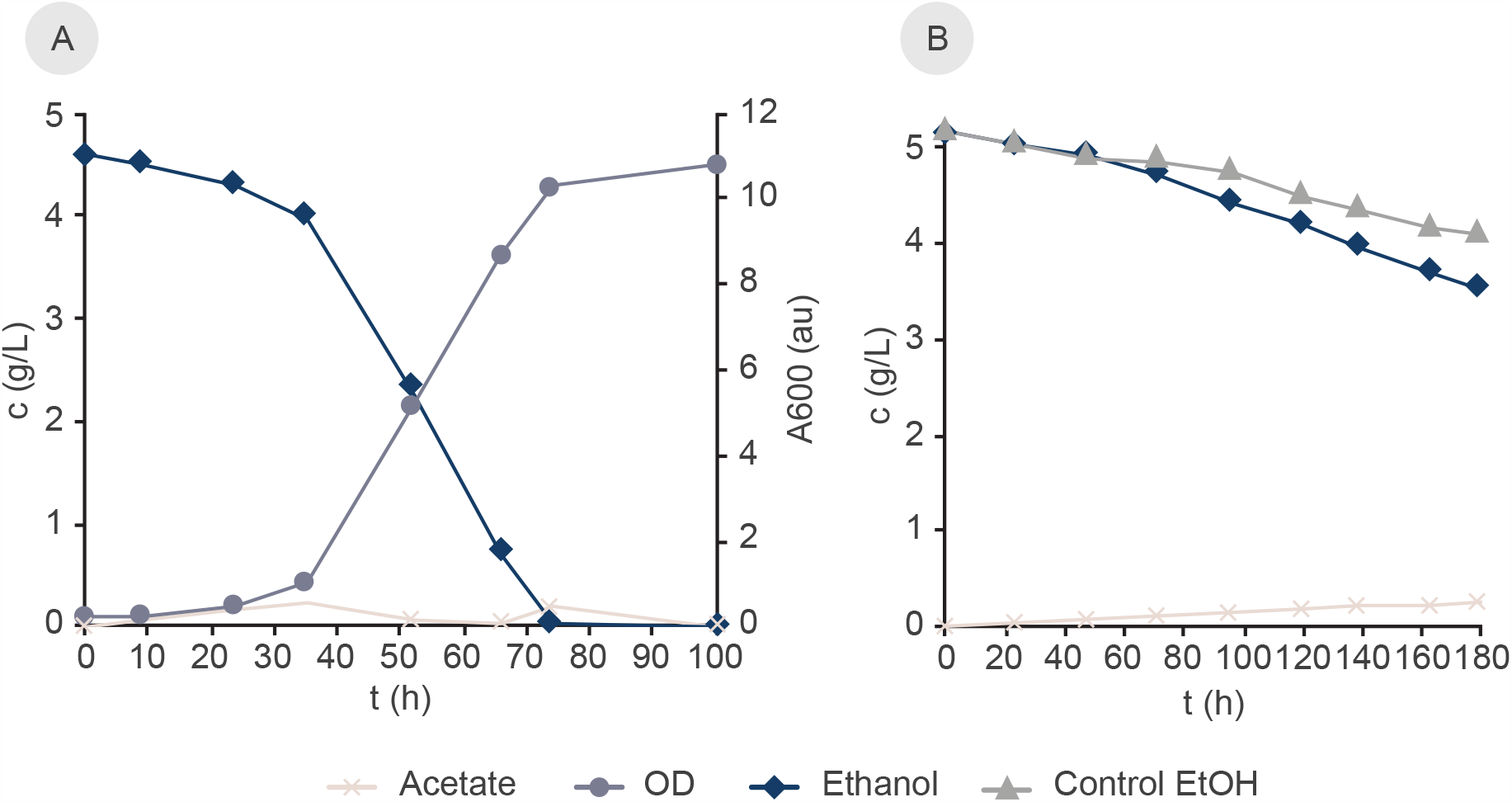
Evaluation of the respiratory metabolism using a defined medium (MM3) with ethanol as the carbon source. Both CP-SCs and CP-LMs were cultured in MM2 for 16 h before being transferred to MM3. (A) CP-SCs (B) CP-LMs. N = 1 each.

In the context of lack of oxygen consumption by CP–LMs, it is noteworthy that the transport of a compound through hydrogels depends on various factors, including the nature of the hydrogel, the compound’s nature and the general circumstances (e.g., temperature). The availability of a compound within a hydrogel can be characterized by flux, which depends on both diffusivity and solubility. The diffusion of most gases like oxygen is several times lower in hydrogels than in water, and carbon dioxide permeation appears to be greater than oxygen but still limited.^41–43^ The evaluation of physical properties and diffusion coefficients of compounds in hydrogels represents an interesting aspect for future research.

Based on our results, we observe that F127-BUM-based LMs are highly restrictive for oxygen transport. They can potentially be developed and deployed as a cost-effective platform to study microaerobic or anaerobic microorganisms, which generally require the creation of an anoxic environment in SCs. An in-depth investigation of oxygen consumption could further help to design oxygen permissible or impermissible LMs based on the desired application. For example, oxygen sensing systems which can be integrated in hydrogels, and could help identifying regions of consumption and the diffusion of oxygen itself within LMs, allowing better 3D printing designs.^43,44^ Nanocomposite hydrogels for the generation of oxygen could further be applied to supply a steady and site-specific release of oxygen.^45^ A mosaic of aerobic and anaerobic regions within a hydrogel could empower the design of new kinds of LMs for multi-kingdom printing using aerobic and anaerobic organisms in parallel.^23^ Nanocomposite hydrogels with organic peroxides for site-specific oxygen release may not be suitable.^46^ Organic peroxides appear to react as photoinitiators and/or create excessive reactive oxygen species, damaging cells in LMs.

### Isolation of cells and flow cytometry

For the metabolism control and efficacy of LMs, it is essential to understand and evaluate cellular phenotypes upon physical confinement. One of the first challenges to understand cellular adaptation in the confined space of crosslinked F127-BUM is the polymeric scaffold itself. Isolating cells to capture the native state of the immobilized cell is necessary to appreciate the molecular basis of phenotypic changes in LMs. For this reason, it is important to avoid harsh treatment protocols to exclude influences on the morphology. We sliced liquid N_2_ quenched LMs, flash-frozen to arrest the cellular processes, with a scalpel and subsequently washed and centrifuged them. We processed the remainder of the sliced pieces in a homogenizer to exfiltrate cells from LMs. Using this approach, we could isolate ca. 23 % of the estimated total cell load, sufficient for cellular characterization and a downstream macromolecular purification (Figure 4A). The homogenizer treatment did not affect the cellular integrity as demonstrated by cell viability assessment (Figure S5). Though, we recognize that further protocol improvements will be necessary for maximizing cell isolation.

**Figure 4:**
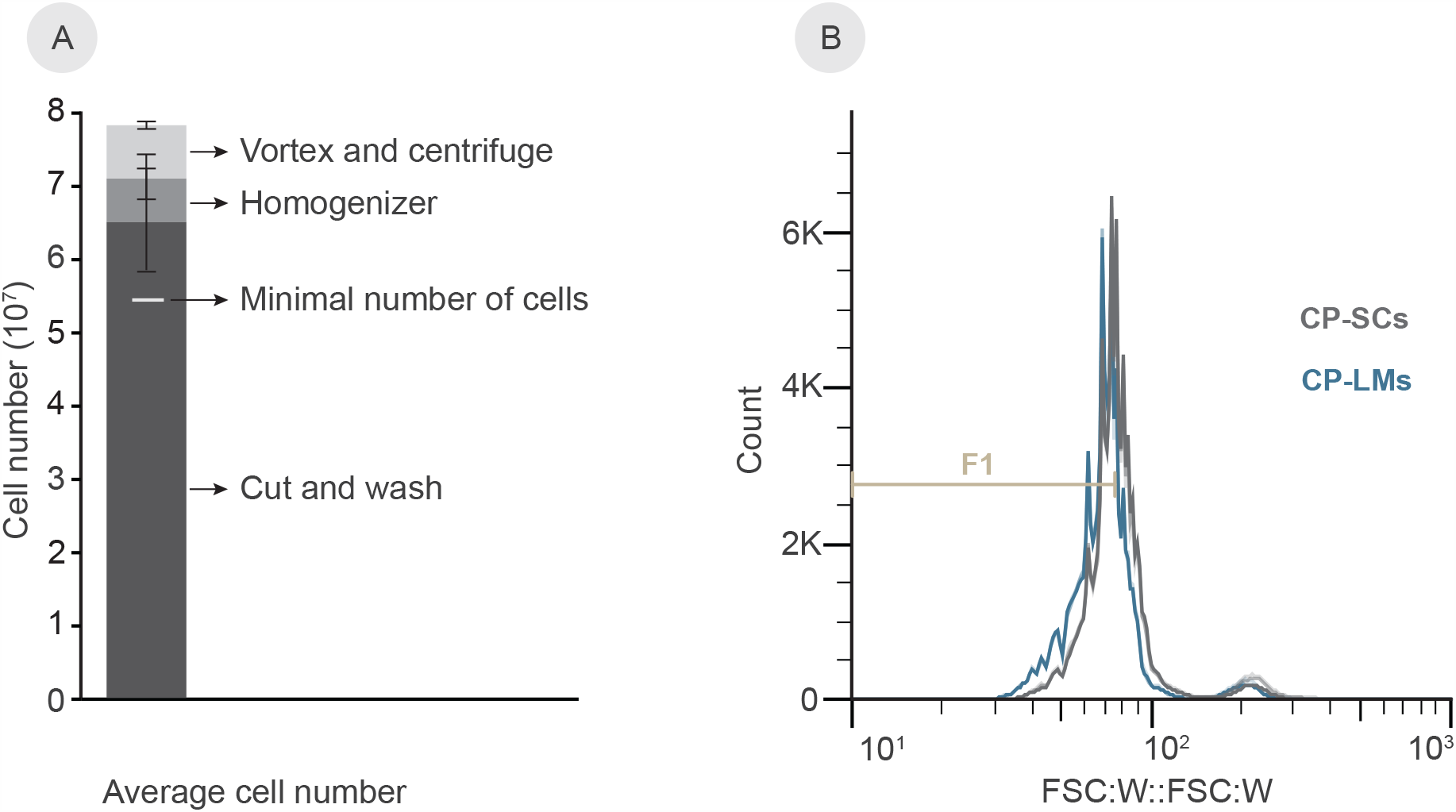
Flow cytometry analysis of isolated cells. (A) Cell isolation from LMs at different processing steps. Minimum indicated as a minimal amount necessary for FC and macromolecule purifications. (B) Forward scatter width histogram with one gate (F1). Gates set to the densest SC area of the first replicate, n = 3 each. All replicates are overlaid and 100,000 cells were acquired for each replicate. Red: CP-SCs, blue: CP-LMs.

We nevertheless tested the macromolecular (DNA, RNA, and proteins) purification from the isolated cells using common protocols. To demonstrate that the genomic DNA amount was sufficient for downstream applications, we performed a PCR amplification of *URA3* (804 bp), a common marker gene, using the purified genomic DNA template (Figure S6A). The genomic DNA could be used either for specific gene amplification or whole-genome sequencing for determining potential mutations during the cultivation, for example, to evaluate the role of UV light or other environmental factors. We assessed the quality of purified RNA molecules using a reverse transcriptase PCR of the most abundant rRNA (26S, terminal amplification 600 bp), and the resulting cDNA was successfully amplified (Figure S6B). We also obtained proteins in sufficient amounts for possible subsequent analyses (data not shown). Our results offer a reproducible method to isolate cells from crosslinked F127 hydrogels enabling further macromolecular investigations. The availability of macromolecules from LMs will allow comparative studies with SCs and other LM systems, e.g., calcium alginate.^34^

FC and fluorescence-activated cell sorting (FACS) are powerful tools to study cells, especially differences of cells under varying conditions, and population heterogeneity. These methodologies allow high throughput analysis relatively quickly, giving reliable and statistically trustworthy results. We assessed immobilized and suspension cells from the glucose-phase for their volume, considering the forward scatter (width) of the cells as a volume proxy, as demonstrated in a former study.^47^ We took formerly immobilized cells from the first isolation round (cut, wash, vortex, centrifuge) for FC, as the homogenizer treatment might influence the membrane thoroughness and thus the volume. The graphs show triplicate layovers for cells obtained from CP–SCs and CP–LMs, where replicates showed high reproducibility (Figure 4B). When comparing the gated populations (small cell fraction) F1, we observed a distinct shift from suspension to immobilized cells. The cells isolated from CP–LM cultures appeared smaller compared to the suspension cells (Figure 4B). Further, we identified a statistical significance in analyzing the fractions in F1 between the conditions, showing higher percentages of CP-LM cells in the small fraction than CP-SCs (Table ST2). The FC results are consistent with the previously reported impact on yeast cell phenotypes upon physical confinement in hydrogels, obtained by analyzing scanning electron microscope images.^20^ The present method offers a more robust cellular phenotyping, with higher throughput readouts, while maintaining a more native cell state. The cell-size differences observed between SCs and CP–LMs are in general agreement with the cell size dependence on growth rate.^48^ Since several factors can influence cell size, including the carbon source, it will be essential to investigate the role of carbon source utilization and biomass conversion in LMs concerning cell size in the future. Carbon source to biomass conversion rate is particularly relevant for developing LMs for 3D printable cell factory applications. The study of immobilized and suspension cells via FACS in the future could be valuable for detecting differently regulated cellular processes, which can be fluorescently labeled.^49^

### Beer fermentation with scaled-up FP-LMs

As we measured higher ethanol yields on glucose and almost no respiration, the application of LMs for ethanol fermentation seemed perfectly suitable. Therefore, we designed a scaled-up structure, as described in the method section. This fabricated structure had a ten-fold higher cell content per gram hydrogel. It contained the beer yeast Bry-97 and fermented sugars in a standard wort formulation for a pale ale type beer. We also designed a classic brewing setup using the same strain in suspension culture, under the same cultivation conditions, including starting sugars used for the FP-LM setting (Figures 5, S7A). The parallel cultivated 3D printed structures remained leakage-free for roughly 5 days, after which cells started to leak from the LMs, forming a suspension culture that slowly settled on the bottom of the beer bottles (Figures 5, S7B). The 3D printed beer appeared more transparent because of a relatively small number of suspension cells compared to the classic setup (Figure S7).

**Figure 5:**
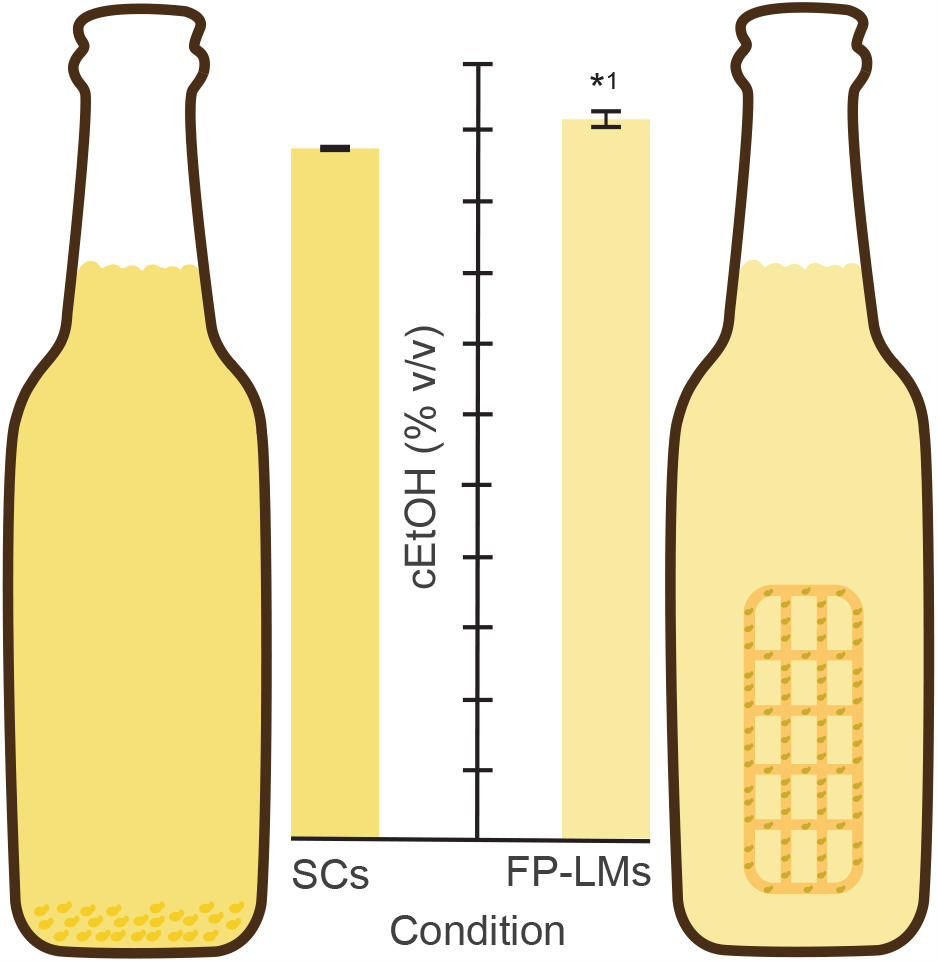
Bottle fermentation with SCs and FP–LMs. Bottle fermentation of a session pale ale style beer. Alcohol content measured after 14 d. ^*,1^ *p* < 0.05 (one tail), significant difference from suspension cells; n = 2, each. Y-axis partitioning: 0.25 %.

Additionally, measuring the beer color revealed differences, with the SC beer having a darker color (Table ST3). After 14 days of brewing, we compared ethanol contents and found a 3.73 % higher alcohol content in the 3D printed beer compared to the classically brewed beer (Figure 5). This ethanol increase in 3D printed beer was significantly higher than classically brewed beer, making 3D printed beer brewing superior in ethanol yield on sugars (one-tailed t-test, Figure 5). The main sugars maltose, glucose and fructose were initially the same for all conditions and replicates, but not detected after 14 d of fermentation. The taste was light in 3D printed beer and appeared sweet with a higher yeast note in SC beer. Both beers were well-carbonated.

Based on our previous cell-retaining experiments and data on the budding yeast, we fabricated the 3D printed structures for the brewery to fully retain the cells. However, we used a common brewer’s yeast strain having different properties, causing cell-leakage after 5 days. Despite the leakage, we demonstrated apparent differences in ethanol yield on sugars, and engineered LMs in the future can be fine-tuned to fully retain the cells during fermentation.^50^ These advances in LMs fabrication will provide additional advantages, such as removing the filtration step after fermentation, and increased savings due to reduced malted barley requirement to achieve the same alcohol level as in classic brewing. Advanced materials and 3D fabrication hold the potential for more cost-effective, sustainable, and reusable microaerobic fermentation systems applicable in brewing.

## Conclusions

Our study offers insights on the metabolism of cells in hydrogels with the potential to control it by creating advanced LMs, permitting the desired cellular phenotype. We successfully demonstrated the control of metabolism in cell-retaining hydrogels whereby immobilized cells were metabolically active in a near-anaerobic environment. As alcohol production currently is the highest volume commercial application in biotechnology, our study demonstrated a proof-of-concept of a 3D printed brewery using yeast in LMs that yielded significantly higher alcohol titers than currently used suspension cultures for beer production. However, due to the lack of information about changes inside of cells in LMs, more investigations are needed to determine the molecular and systems-level changes that happen as the cellular phenotype alters upon physical encapsulation. We present a method for cell isolation that will be helpful for such investigations in the future. Further technological advances using LMs could be envisioned to help in sustainable wastewater treatment, fighting against global warming solutions by allowing the cultivation of anaerobic cultures such as acetogens and methanogens, which utilize CO_2_ or CH_4_ as a source of energy and biomass formation.

## Acknowledgments

This work received funding from the European Union’s Horizon 2020 research and innovation program under grant agreement No. 668997, and the Estonian Research Council (grant PUT1488P). TB additionally acknowledges mobility grants MB-2018-1/25 by Bayerisches Hochschulzentrum für Mittel-, Ost-und Südosteuropa (BAYHOST) and Erasmus+ (European Union). AN acknowledges the National Science Foundation (Grant No. 1752972), UW CoMotion Innovation Grant, and UW Royalty Research Fund for financial support of this work. We thank Laura Kibena for excellent technical assistance, Aleksandr Illarionov, Lorena Lima and Triinu Visnapuu for providing reagents. Thanks to Dr. Nemailla Bonturi for an insightful discussion.

## Conflict of Interest

None

## Notes

### Competing Interest Statement

The authors have declared no competing interest.

## References

(1) Hull, C. W. Apparatus for Production of Three-Dimensional Objects by Stereolithography. Patent nr. 4,575,330, August 8, 1984.

(2) Klebe, R. J. Cytoscribing: A Method for Micropositioning Cells and the Construction of Two- and Three-Dimensional Synthetic Tissues. Exp. Cell Res. 1988, 179 (2), 362–373. https://doi.org/10.1016/0014-4827(88)90275-3.

(3) Jose, R. R.; Rodriguez, M. J.; Dixon, T. A.; Omenetto, F.; Kaplan, D. L. Evolution of Bioinks and Additive Manufacturing Technologies for 3D Bioprinting. ACS Biomater. Sci. Eng. 2016, 2 (10), 1662–1678. https://doi.org/10.1021/acsbiomaterials.6b00088.

(4) Peng, W.; Unutmaz, D.; Ozbolat, I. T. Bioprinting towards Physiologically Relevant Tissue Models for Pharmaceutics. Trends in Biotechnology. Elsevier Ltd September 1, 2016, pp 722–732. https://doi.org/10.1016/j.tibtech.2016.05.013.

(5) Das, S.; Basu, B. An Overview of Hydrogel-Based Bioinks for 3D Bioprinting of Soft Tissues. Journal of the Indian Institute of Science. Springer International Publishing September 1, 2019, pp 405–428. https://doi.org/10.1007/s41745-019-00129-5.

(6) Saygili, E.; Dogan-Gurbuz, A. A.; Yesil-Celiktas, O.; Draz, M. S. 3D Bioprinting: A Powerful Tool to Leverage Tissue Engineering and Microbial Systems. Bioprinting. Elsevier B.V. June 1, 2020, p e00071. https://doi.org/10.1016/j.bprint.2019.e00071.

(7) Meinert, C.; Schrobback, K.; Hutmacher, D. W.; Klein, T. J. A Novel Bioreactor System for Biaxial Mechanical Loading Enhances the Properties of Tissue-Engineered Human Cartilage. Sci. Rep. 2017, 7 (1), 1–14. https://doi.org/10.1038/s41598-017-16523-x.

(8) Nichol, J. W.; Khademhosseini, A. Modular Tissue Engineering: Engineering Biological Tissues from the Bottom Up. Soft Matter 2009, 5 (7), 1312–1319. https://doi.org/10.1039/b814285h.

(9) Kyle, S. 3D Printing of Bacteria: The Next Frontier in Biofabrication. Trends in Biotechnology. Elsevier Ltd April 1, 2018, pp 340–341. https://doi.org/10.1016/j.tibtech.2018.01.010.

(10) Balasubramanian, S.; Aubin-Tam, M. E.; Meyer, A. S. 3D Printing for the Fabrication of Biofilm-Based Functional Living Materials. ACS Synthetic Biology. American Chemical Society July 19, 2019, pp 1564–1567. https://doi.org/10.1021/acssynbio.9b00192.

(11) Huang, J.; Liu, S.; Zhang, C.; Wang, X.; Pu, J.; Ba, F.; Xue, S.; Ye, H.; Zhao, T.; Li, K.; Wang, Y.; Zhang, J.; Wang, L.; Fan, C.; Lu, T. K.; Zhong, C. Programmable and Printable Bacillus Subtilis Biofilms as Engineered Living Materials. Nat. Chem. Biol. 2019, 15 (1), 34–41. https://doi.org/10.1038/s41589-018-0169-2.

(12) Landreau, M.; Byson, S. J.; You, H. J.; Stahl, D. A.; Winkler, M. K. H. Effective Nitrogen Removal from Ammonium-Depleted Wastewater by Partial Nitritation and Anammox Immobilized in Granular and Thin Layer Gel Carriers. Water Res. 2020, 183, 116078. https://doi.org/10.1016/j.watres.2020.116078.

(13) Gilbert, C.; Tang, T.-C.; Ott, W.; Dorr, B. A.; Shaw, W. M.; Sun, G. L.; Lu, T. K.; Ellis, T. Living Materials with Programmable Functionalities Grown from Engineered Microbial Co-Cultures. Nat. Mater. 2021, 1–10. https://doi.org/10.1038/s41563-020-00857-5.

(14) Shachaf, Y.; Gonen-Wadmany, M.; Seliktar, D. The Biocompatibility of Pluronic®F127 Fibrinogen-Based Hydrogels. Biomaterials 2010, 31 (10), 2836–2847. https://doi.org/10.1016/J.BIOMATERIALS.2009.12.050.

(15) Wu, C.-J.; Gaharwar, A. K.; Chan, B. K.; Schmidt, G. Mechanically Tough Pluronic F127/Laponite Nanocomposite Hydrogels from Covalently and Physically Cross-Linked Networks. Macromolecules 2011, 44 (20), 8215–8224. https://doi.org/10.1021/ma200562k.

(16) Gioffredi, E.; Boffito, M.; Calzone, S.; Maria Giannitelli, S.; Rainer, A.; Trombetta, M.; Mozetic, P.; Chiono, V. Pluronic F127 Hydrogel Characterization and Biofabrication in Cellularized Constructs for Tissue Engineering Applications. Procedia CIRP 2016, 49, 125–132. https://doi.org/10.1016/j.procir.2015.11.001.

(17) Saha, A.; Johnston, T. G.; Shafranek, R. T.; Goodman, C. J.; Zalatan, J. G.; Storti, D. W.; Ganter, M. A.; Nelson, A. Additive Manufacturing of Catalytically Active Living Materials. ACS Appl. Mater. Interfaces 2018, acsami.8b02719. https://doi.org/10.1021/acsami.8b02719.

(18) Millik, S. C.; Dostie, A. M.; Karis, D. G.; Smith, P. T.; McKenna, M.; Chan, N.; Curtis, C. D.; Nance, E.; Theberge, A. B.; Nelson, A. 3D Printed Coaxial Nozzles for the Extrusion of Hydrogel Tubes toward Modeling Vascular Endothelium. Biofabrication 2019, 11 (4), 045009. https://doi.org/10.1088/1758-5090/ab2b4d.

(19) Johnston, T. G.; Yuan, S.-F.; Wagner, J. M.; Yi, X.; Saha, A.; Smith, P.; Nelson, A.; Alper, H. S. Compartmentalized Microbes and Co-Cultures in Hydrogels for on-Demand Bioproduction and Preservation. Nat. Commun. 2020, 11 (1), 563. https://doi.org/10.1038/s41467-020-14371-4.

(20) Priks, H.; Butelmann, T.; Illarionov, A.; Johnston, T. G.; Fellin, C.; Tamm, T.; Nelson, A.; Kumar, R.; Lahtvee, P.-J. Physical Confinement Impacts Cellular Phenotype within Living Materials. ACS Appl. Bio Mater.2020. https://doi.org/10.1021/acsabm.0c00335.

(21) Boydston, A. J.; Cao, B.; Nelson, A.; Ono, R. J.; Saha, A.; Schwartz, J. J.; Thrasher, C. J. Additive Manufacturing with Stimuli-Responsive Materials. Journal of Materials Chemistry Royal Society of Chemistry 2018, pp 20621–20645. https://doi.org/10.1039/C8TA07716A.

(22) Smith, P. T.; Basu, A.; Saha, A.; Nelson, A. Chemical Modification and Printability of Shear-Thinning Hydrogel Inks for Direct-Write 3D Printing. Polym. (United Kingdom) 2018, 1–9. https://doi.org/10.1016/j.polymer.2018.01.070.

(23) Johnston, T. G.; Fillman, J. P.; Priks, H.; Butelmann, T.; Tamm, T.; Kumar, R.; Lahtvee, P.-J.; Nelson, A. Cell-Laden Hydrogels for Multikingdom 3D Printing. Macromol. Biosci. 2020, 2000121. https://doi.org/10.1002/mabi.202000121.

(24) Cushing, M. C.; Anseth, K. S. Hydrogel Cell Cultures. Science. American Association for the Advancement of Science May 25, 2007, pp 1133–1134. https://doi.org/10.1126/science.1140171.

(25) Rademakers, T.; Horvath, J. M.; van Blitterswijk, C. A.; LaPointe, V. L. S. Oxygen and Nutrient Delivery in Tissue Engineering: Approaches to Graft Vascularization. Journal of Tissue Engineering and Regenerative Medicine. John Wiley and Sons Ltd October 1, 2019, pp 1815–1829. https://doi.org/10.1002/term.2932.

(26) Heffernan, J. K.; Valgepea, K.; de Souza Pinto Lemgruber, R.; Casini, I.; Plan, M.; Tappel, R.; Simpson, S. D.; Köpke, M.; Nielsen, L. K.; Marcellin, E. Enhancing CO2-Valorization Using Clostridium Autoethanogenum for Sustainable Fuel and Chemicals Production. Front. Bioeng. Biotechnol. 2020, 8, 204. https://doi.org/10.3389/fbioe.2020.00204.

(27) Kim, E. S.; Kim, B. S.; Kim, K. Y.; Woo, H. M.; Lee, S. M.; Um, Y. Aerobic and Anaerobic Cellulose Utilization by Paenibacillus Sp. CAA11 and Enhancement of Its Cellulolytic Ability by Expressing a Heterologous Endoglucanase. J. Biotechnol. 2018, 268, 21–27. https://doi.org/10.1016/j.jbiotec.2018.01.007.

(28) Lee, D.; Lloyd, N. D. R.; Pretorius, I. S.; Borneman, A. R. Heterologous Production of Raspberry Ketone in the Wine Yeast Saccharomyces Cerevisiae via Pathway Engineering and Synthetic Enzyme Fusion. Microb. Cell Fact. 2016, 15 (1), 49. https://doi.org/10.1186/s12934-016-0446-2.

(29) Kumar, R.; Lahtvee, P.-J. Proteome Overabundance Enables Respiration but Limitation Onsets Carbon Overflow. bioRxiv 2020, 2020.02.20.957662. https://doi.org/10.1101/2020.02.20.957662.

(30) Avalos, J. L.; Fink, G. R.; Stephanopoulos, G. Compartmentalization of Metabolic Pathways in Yeast Mitochondria Improves the Production of Branched-Chain Alcohols. Nat. Biotechnol. 2013, 31 (4), 335–341. https://doi.org/10.1038/nbt.2509.

(31) Valgepea, K.; de Souza Pinto Lemgruber, R.; Meaghan, K.; Palfreyman, R. W.; Abdalla, T.; Heijstra, B. D.; Behrendorff, J. B.; Tappel, R.; Köpke, M.; Simpson, S. D.; Nielsen, L. K.; Marcellin, E. Maintenance of ATP Homeostasis Triggers Metabolic Shifts in Gas-Fermenting Acetogens. Cell Syst. 2017, 4 (5), 505-515.e5. https://doi.org/10.1016/j.cels.2017.04.008.

(32) Shaw, D. R.; Ali, M.; Katuri, K. P.; Gralnick, J. A.; Reimann, J.; Mesman, R.; van Niftrik, L.; Jetten, M. S. M.; Saikaly, P. E. Extracellular Electron Transfer-Dependent Anaerobic Oxidation of Ammonium by Anammox Bacteria. Nat. Commun. 2020, 11 (1), 1–12. https://doi.org/10.1038/s41467-020-16016-y.

(33) Zekker, I.; Raudkivi, M.; Artemchuk, O.; Rikmann, E.; Priks, H.; Jaagura, M.; Tenno, T. Mainstream-Sidestream Wastewater Switching Promotes Anammox Nitrogen Removal Rate in Organic-Rich, Low-Temperature Streams. Environ. Technol. (United Kingdom) 2020. https://doi.org/10.1080/09593330.2020.1721566.

(34) Nagarajan, S.; Kruckeberg, A. L.; Schmidt, K. H.; Kroll, E.; Hamilton, M.; McInnerney, K.; Summers, R.; Taylor, T.; Rosenzweig, F. Uncoupling Reproduction from Metabolism Extends Chronological Lifespan in Yeast. Proc. Natl. Acad. Sci. U. S. A. 2014, 111 (15), E1538–47. https://doi.org/10.1073/pnas.1323918111.

(35) Lõoke, M.; Kristjuhan, K.; Kristjuhan, A. Extraction of Genomic DNA from Yeasts for PCR-Based Applications. Biotechniques 2011, 50 (5), 325–328. https://doi.org/10.2144/000113672.

(36) Sherman, F. Getting Started with Yeast. Methods Enzymol. 2002, 350, 3–41. https://doi.org/10.1016/S0076-6879(02)50954-X.

(37) Klis, F. M.; de Koster, C. G.; Brul, S. Cell Wall-Related Bionumbers and Bioestimates of Saccharomyces Cerevisiae and Candida Albicans. Eukaryot. Cell 2014, 13 (1), 2–9. https://doi.org/10.1128/EC.00250-13.

(38) Li, S. C.; Kane, P. M. The Yeast Lysosome-like Vacuole: Endpoint and Crossroads. Biochimica et Biophysica Acta - Molecular Cell Research. Elsevier April 1, 2009, pp 650–663. https://doi.org/10.1016/j.bbamcr.2008.08.003.

(39) Illarionov, A.; Lahtvee, P.-J.; Kumar, R. Characterization of Potassium, Sodium and Their Interactions Effects in Yeasts. bioRxiv 2020, 2020.10.22.350355. https://doi.org/10.1101/2020.10.22.350355.

(40) Vanoni, M.; Rossi, R. L.; Querin, L.; Zinzalla, V.; Alberghina, L. Glucose Modulation of Cell Size in Yeast. In Biochemical Society Transactions; Portland Press, 2005; Vol. 33, pp 294–296. https://doi.org/10.1042/BST0330294.

(41) Refojo, M. F. Mechanism of Gas Transport through Contact Lenses. J. Am. Optom. Assoc. 1979, 50 (3), 285–287.

(42) Holden, B. A.; Ross, R.; Jenkins, J. Hydrogel Contact Lenses Impede Carbon Dioxide Efflux from the Human Cornea. Curr. Eye Res. 1987, 6 (11), 1283–1290. https://doi.org/10.3109/02713688708997553.

(43) Figueiredo, L.; Pace, R.; D’Arros, C.; Réthoré, G.; Guicheux, J.; Le Visage, C.; Weiss, P. Assessing Glucose and Oxygen Diffusion in Hydrogels for the Rational Design of 3D Stem Cell Scaffolds in Regenerative Medicine. J. Tissue Eng. Regen. Med. 2018, 12 (5), 1238–1246. https://doi.org/10.1002/term.2656.

(44) Trampe, E.; Koren, K.; Akkineni, A. R.; Senwitz, C.; Krujatz, F.; Lode, A.; Gelinsky, M.; Kühl, M. Functionalized Bioink with Optical Sensor Nanoparticles for O2 Imaging in 3D-Bioprinted Constructs. Adv. Funct. Mater. 2018, 28 (45), 1804411. https://doi.org/10.1002/adfm.201804411.

(45) Camci-Unal, G.; Alemdar, N.; Annabi, N.; Khademhosseini, A. Oxygen-Releasing Biomaterials for Tissue Engineering. Polym. Int. 2013, 62 (6), 843–848. https://doi.org/10.1002/pi.4502.

(46) Kehr, N. S.; Motealleh, A. Injectable Oxygen-Generating Nanocomposite Hydrogels with Prolonged Oxygen Delivery for Enhanced Cell Proliferation under Hypoxic and Normoxic Conditions. J. Mater. Chem. B 2020. https://doi.org/10.1039/D0TB00885K.

(47) Tzur, A.; Moore, J. K.; Jorgensen, P.; Shapiro, H. M.; Kirschner, M. W. Optimizing Optical Flow Cytometry for Cell Volume-Based Sorting and Analysis. PLoS One 2011, 6 (1), e16053. https://doi.org/10.1371/journal.pone.0016053.

(48) Aldea, M.; Jenkins, K.; Csikász-Nagy, A. Growth Rate as a Direct Regulator of the Start Network to Set Cell Size. Front. Cell Dev. Biol. 2017, 5 (MAY), 57. https://doi.org/10.3389/fcell.2017.00057.

(49) Haase, S. B.; Reed, S. I. Improved Flow Cytometric Analysis of the Budding Yeast Cell Cycle. Cell Cycle 2002, 1 (2), 117–121. https://doi.org/10.4161/cc.1.2.114.

(50) Guo, S.; Dubuc, E.; Rave, Y.; Verhagen, M.; Twisk, S. A. E.; Van Der Hek, T.; Oerlemans, G. J. M.; Van Den Oetelaar, M. C. M.; Van Hazendonk, L. S.; Brüls, M.; Eijkens, B. V.; Joostens, P. L.; Keij, S. R.; Xing, W.; Nijs, M.; Stalpers, J.; Sharma, M.; Gerth, M.; Boonen, R. J. E. A.; Verduin, K.; Merkx, M.; Voets, I. K.; De Greef, T. F. A. Engineered Living Materials Based on Adhesin-Mediated Trapping of Programmable Cells. ACS Synth. Biol. 2020, 9 (3), 475–485. https://doi.org/10.1021/acssynbio.9b00404.

