## Supplemental Figures (S) and Tables (ST) for "Metabolism Control in 3D Printed Living Materials"

### Affiliations

### Figures (S)

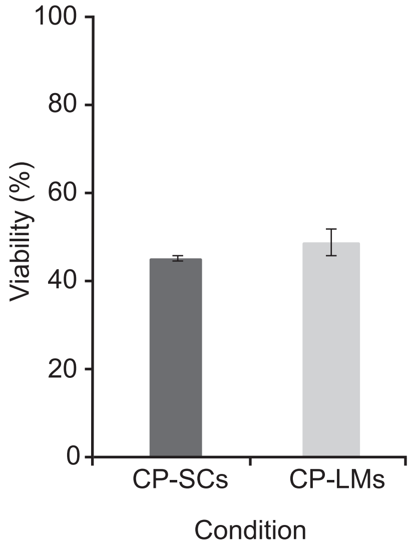


**Figure S1: Viability of CP-SCs and CP-LMs after stepwise thawing.**

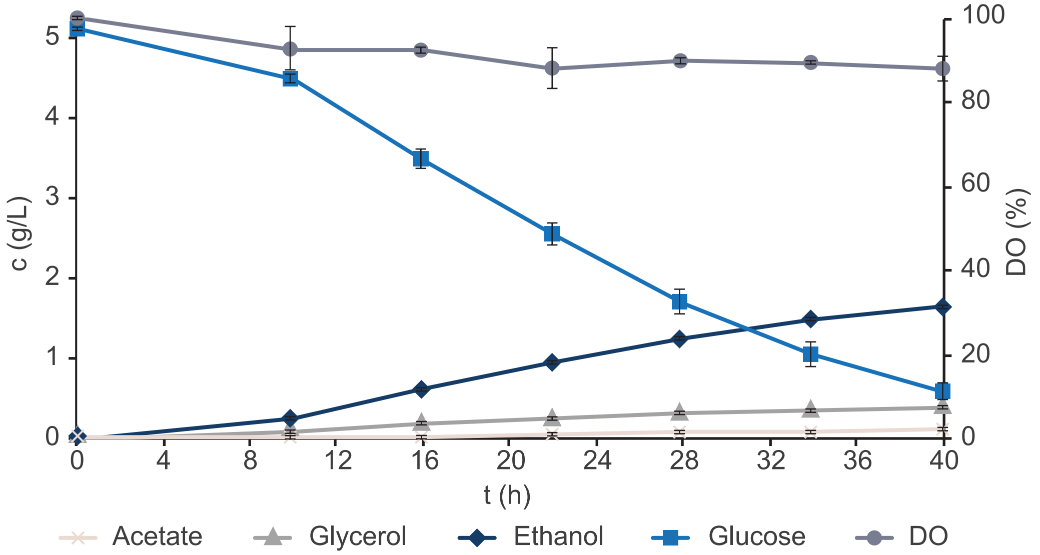


**Figure S2: Yeast physiology in glucose phases of batch cultivations in FP-LMs.** DO: dissolved oxygen. Error bars represent standard deviations, n = 3.

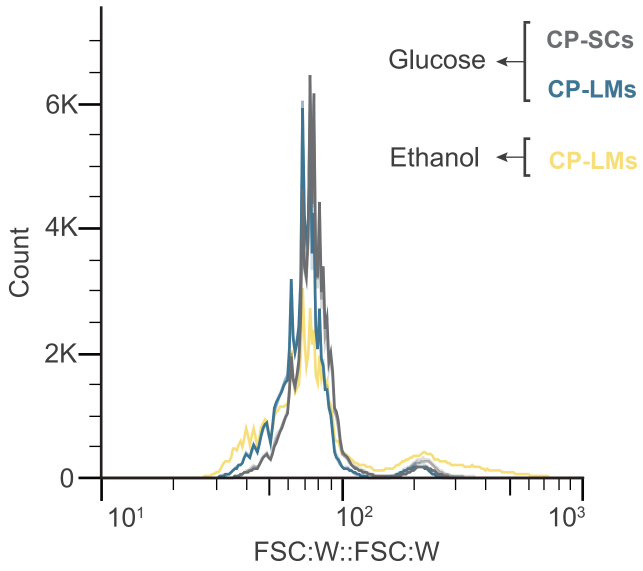


**Figure S3: Flow cytometry of glucose and ethanol phase.** Glucose plots are the same as in Figure 4B. The yellow plot represents CP-LM cells in the ethanol phase. Forward scatter width histogram, 100,000 cells were acquired for each replicate.

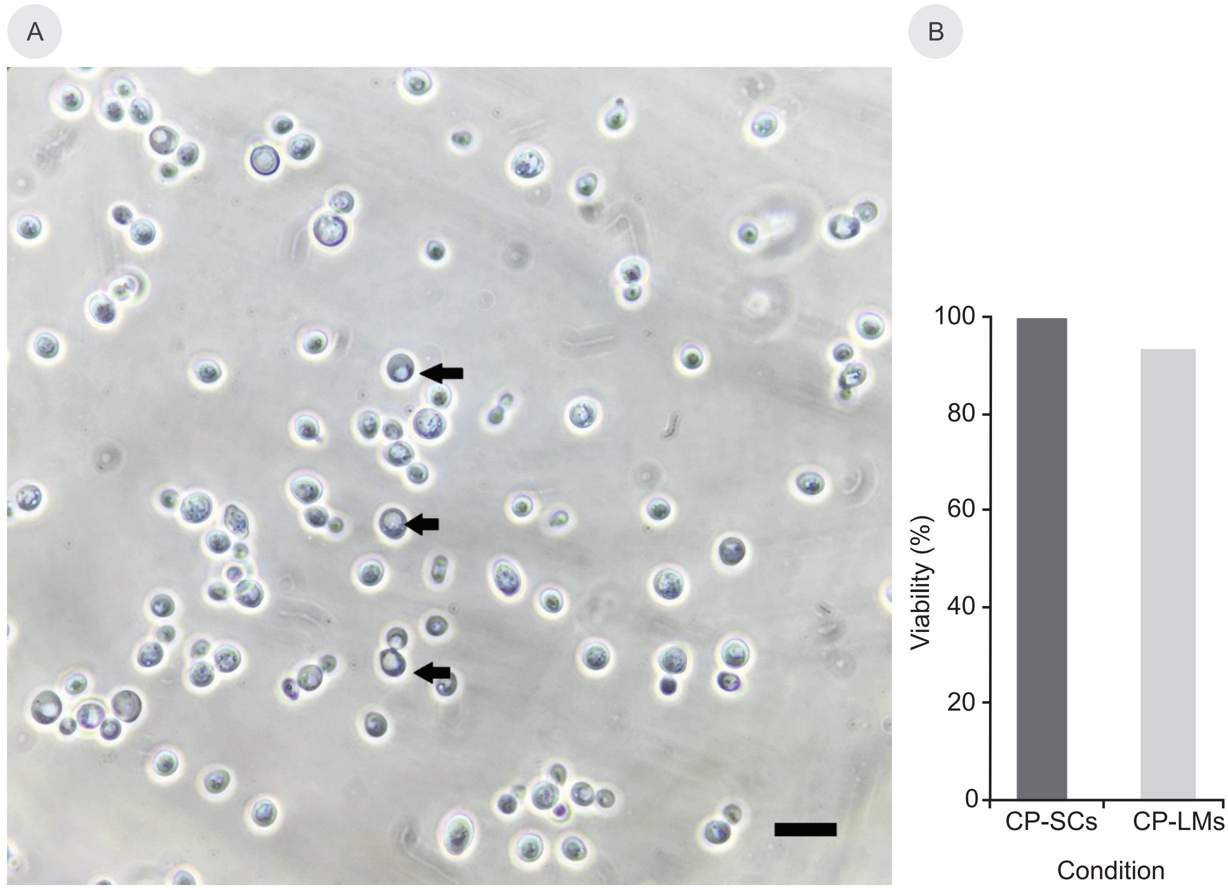

**Figure S4: Stress during ethanol cultivation in CP-LMs.** (A) Increased vacuole size in yeast cells with ethanol as a carbon source, indicated by black arrows. Scale bar 10 µm. (B) Viability of cells in CP-LM and CP-SCs after cultivation.

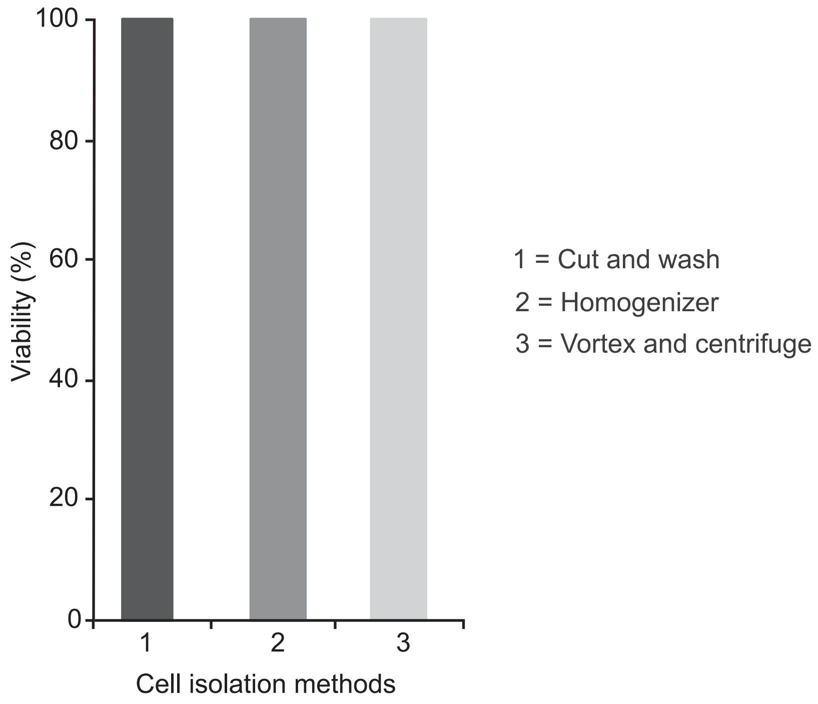


**Figure S5: Viability of yeast cells from LMs after different isolation methods.**

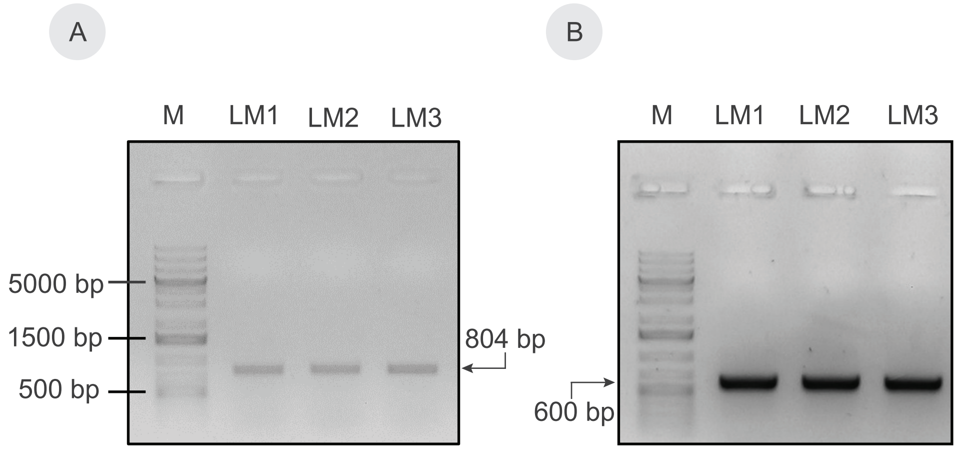

**Figure S6: Agarose gel images of amplified nucleic acids.** (A) 1 % agarose gel with amplified *URA3* from genomic DNA. (B) 1 % agarose gel with amplified 26S cDNA from purified and reverse transcribed RNA. M; marker.

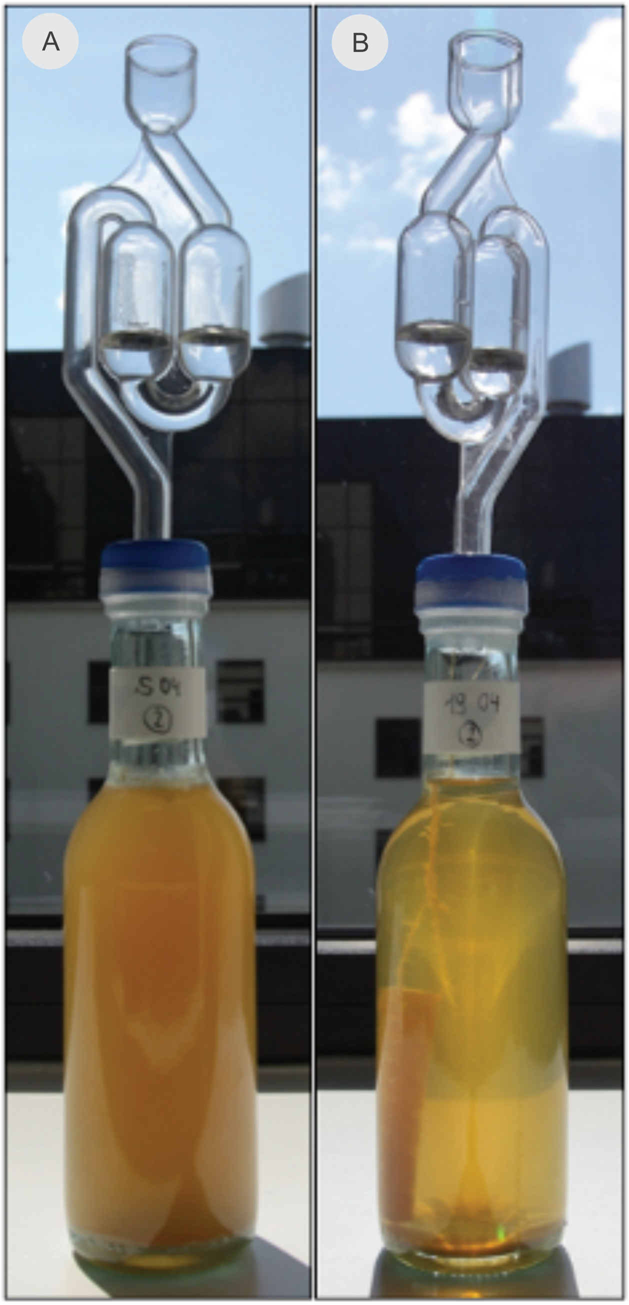


**Figure S7: Images of bottle-fermented pale ale style beers by SCs and FP–LM.** (A) SCs, (B) FP-LM.

**Tables (ST)**

**Table ST1. Yields on glucose (G) in FP-LMs.** Calculation done over the complete time span. ** $p < 0.01$, significant difference from SCs in Table 2. n = 3.

| | $Y_{G,glycerol}$ (g/g) | $Y_{G,acetate}$ (g/g) | $Y_{G,ethanol}$ (g/g) |
|---|---|---|---|
| FP-LMs (Mean ±Std) | 0.0900 ±0.00 ** | 0.0166 ±0.00 ** | 0.4593 ±0.02 ** |

**Table ST2: Percentage of cells in small cell fractions.** Table corresponds to fractions mentioned in Figure 4B. Right column: mean ±Std. ** $p < 0.01$, significant difference from SCs, n = 3.

| | | | |
|---|---|---|---|
| CP-SCs | F1 SC1 | 52.5 % | 50.5 ± 2.3 % |
| | F1 SC2 | 51.7 % | |
| | F1 SC3 | 47.3 % | |
| CP-LMs | F1 LM1 | 63.1 % | 63.4 ± 0.2 % ** |
| | F1 LM2 | 63.7 % | |
| | F1 LM3 | 63.5 % | |

**Table ST3: Beer color according to different methods.** $A_{430}$, Standard Reference Method (SRM), European Brewery Convention (EBC), n = 2 each. The color is typical for a pale ale style beer.

| | **$A_{430}$ (au)** | **SRM (au)** | **EBC (au)** |
|---|---|---|---|
| **SCs** | 0.377 (±0.005) | 4.788 (±0.064) | 9.425 (±0.125) |
| **FP-LMs** | 0.358 (±0) | 4.547 (±0) | 8.950 (±0) |
